# Platelet bioenergetics correlate with skeletal muscle metabolism in C57BL/6J mice

**DOI:** 10.1101/2024.12.03.626404

**Authors:** Mia S Wilkinson, Emily J Ferguson, Justin Bureau, Jennifer LMH Veeneman, Patricia DA Lima, Chris McGlory, Kimberly J Dunham-Snary

## Abstract

Skeletal muscle insulin resistance is a key step in progression of cardiometabolic disease, and impaired mitochondrial bioenergetics has been implicated. However, mitochondrial bioenergetic research in skeletal muscle is limited by the need for muscle biopsies. We sought to determine if platelet bioenergetics could be used as a minimally invasive surrogate for skeletal muscle bioenergetics. Multiple parameters of mitochondrial respiration, measured by high resolution respirometry, correlated between platelets and gastrocnemius muscle in mice. We propose the coupling state of platelet mitochondria reflects that of skeletal muscle in mice, providing a foundation for future research on using platelets as a liquid biopsy for muscle mitochondrial health in cardiometabolic disease, offering early insights into muscle metabolism to enhance clinical biomarker implementation.

**Graphical Abstract:** 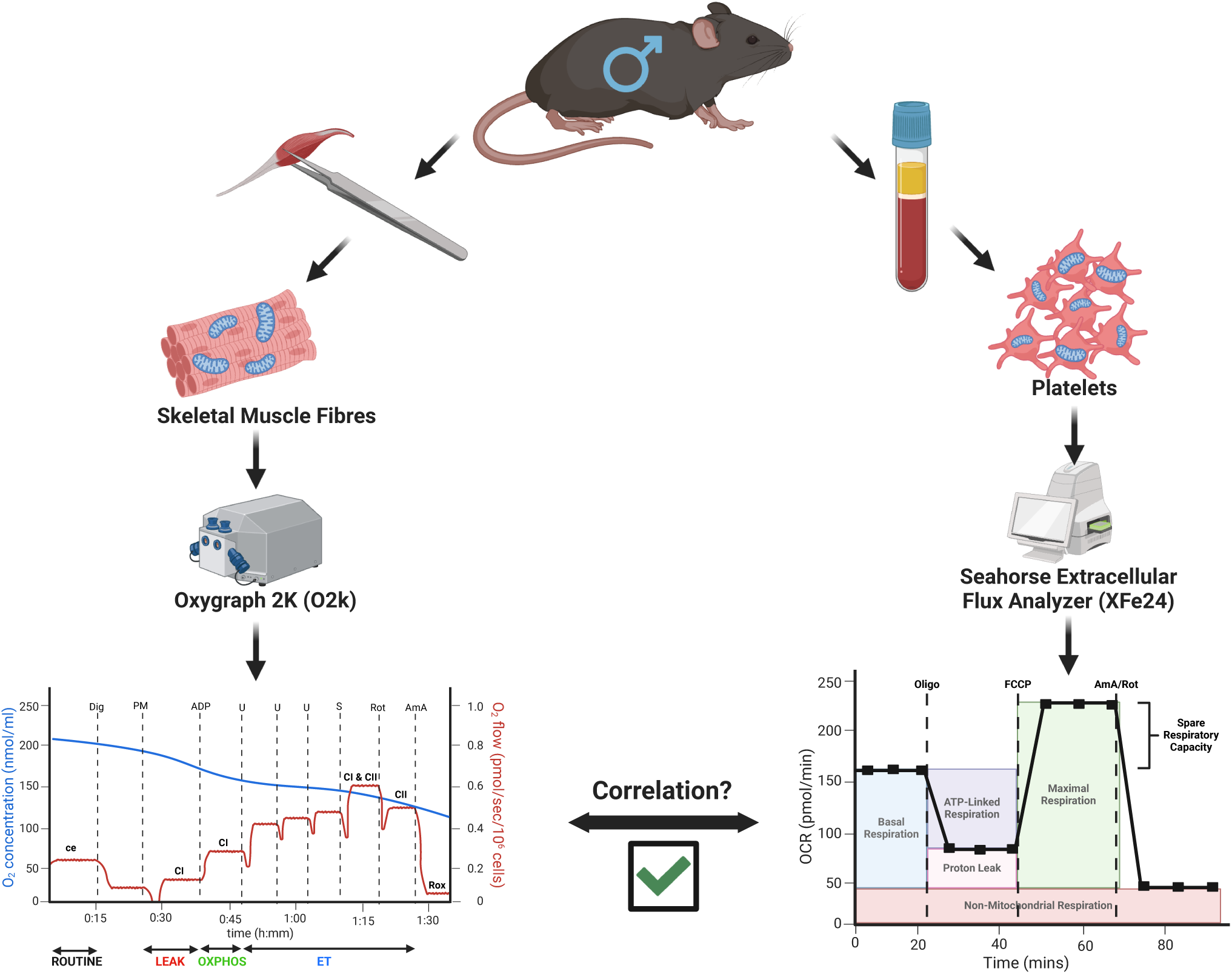

## Introduction

Cardiometabolic disease (CMD) is highly prevalent worldwide, with obesity alone responsible for an ***annual*** healthcare cost of $173 billion in the United States^1–3^. CMD, or metabolic syndrome, is diagnosed by the ≥3 of the following factors: abdominal obesity (population-/country-specific definitions); elevated triglycerides (≥150 mg/dL); reduced HDL-c (<40 mg/dL, males; <50 mg/dL, females); hypertension (systolic ≥130 and/or diastolic ≥85 mmHg); elevated fasting glucose (≥100 mg/dL)^4^. This pre-disease state increases cardiovascular disease (CVD) and Type II diabetes (T2DM) risk, leading causes morbidity and mortality worldwide, with heart diseases accounting for one third of all deaths, globally^5^. Rising CMD incidence highlights the insufficiency of diet, exercise, and current treatments to address underlying genetic and biomolecular factors impacting disease etiology^6^. A critical step in the progression from CMD to CVD and T2DM is the emergence of insulin resistance, resulting in metabolic disturbances and compromised tissue function^7,8^. Skeletal muscle is the largest insulin-sensitive tissue in the body and exhibits functional decline during progression of CMD, CVD, and T2DM^7–10^. Mitochondria, a hub of cellular metabolism, play a fundamental role in CMD-related skeletal muscle insulin resistance and metabolic inflexibility through reduced oxidative capacity, increased reactive oxygen species (ROS), and impaired coupling^8,11–13^.

The study of skeletal muscle mitochondrial respiration has emerged as a translational tool for characterization of mitochondrial function in human pathology^14–16^. In this way, skeletal muscle mitochondria serve as the ‘canary in a coal mine’ to provide early warning of bioenergetic crisis for determining severity and progression of complex and multifactorial disease like CMD. While skeletal muscle mitochondrial bioenergetic dysfunction is recognized as a major player in age- and disease-associated decline in skeletal muscle health^13,17,18^, both a unifying mechanism and targeted treatments are lacking. This is, in part, due to the requirement of a skeletal muscle biopsy for bioenergetic assessment, which are invasive, costly, and often contraindicated^19^. These factors, combined with unpredictable patient compliance and hesitation, create a bottleneck for research, hindering characterization of pathology-associated mitochondrial dysfunction and limiting the clinical utility of tissue bioenergetics as a biomarker. The emergence of platelet bioenergetics as a surrogate for skeletal muscle mitochondrial function combats the aforementioned barriers^20–22^. Indeed, peripheral blood platelets, the cell fragments of megakaryocytes, have been shown to reflect skeletal muscle mitochondrial metabolism in animal models and humans^23–25^. The comparative ease of obtaining platelets, their abundance, highly metabolic phenotype, and high responsiveness to environmental changes all support their candidacy as a biomarker of mitochondrial function^19^. Platelet isolation from whole blood is quick, economic, and does not necessitate specialized equipment or training^26,27^. Also, lack of a nucleus makes platelets an ideal tool for investigating the contribution of mitochondrial DNA to mitochondrial function^28^.

The comparative inaccessibility of skeletal muscle mitochondria, coupled with their status as a master regulator of tissue health, necessitates an accessible and reliable method of assessment^19^. The translational potential of platelet bioenergetics is being explored across disease states; its suitability as a biomarker for early metabolic changes in CMD has not been explored. Due to limited to control in human metabolic studies (i.e. high variability in lifestyle factors), murine models are commonly used to model metabolic disease caused by nutrient overload or manipulation of identified genetic factors. In this proof-of-concept study, we sought to assess if correlation between platelet and skeletal muscle bioenergetics demonstrated in other disease models holds true in mice. Establishing correlation in healthy mice allows for future studies investigating metabolic disease in preclinical models, for eventual translation to human participants; this method represents a promising ‘liquid biopsy’ that may offer early insights into muscle metabolic health in CMD and improve prognostic and diagnostic biomarker implementation in clinical practice. Using male C57BL/6J mice, a strain commonly utilized for preclinical obesity studies, we assessed whether platelet bioenergetics could serve as a surrogate measure for skeletal muscle energetics in healthy mice.

## STAR★Methods

### Key resources table

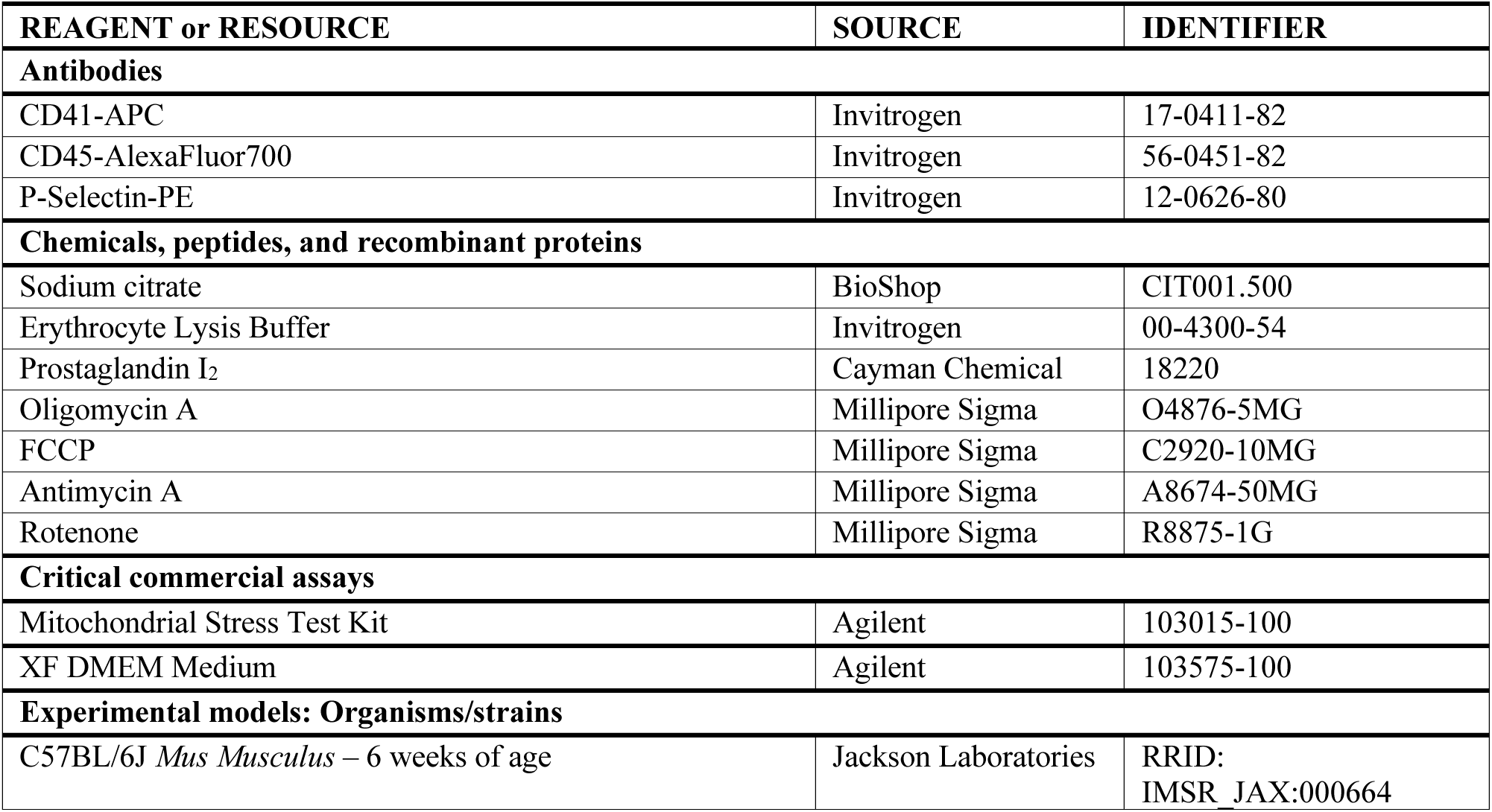

| REAGENT or RESOURCE | SOURCE | IDENTIFIER |
| --- | --- | --- |
| **Antibodies** |  |  |
| CD41-APC | Invitrogen | 17-0411-82 |
| CD45-AlexaFluor700 | Invitrogen | 56-0451-82 |
| P-Selectin-PE | Invitrogen | 12-0626-80 |
| **Chemicals, peptides, and recombinant proteins** |  |  |
| Sodium citrate | BioShop | CIT001.500 |
| Erythrocyte Lysis Buffer | Invitrogen | 00-4300-54 |
| Prostaglandin I<sub>2</sub> | Cayman Chemical | 18220 |
| Oligomycin A | Millipore Sigma | O4876-5MG |
| FCCP | Millipore Sigma | C2920-10MG |
| Antimycin A | Millipore Sigma | A8674-50MG |
| Rotenone | Millipore Sigma | R8875-1G |
| **Critical commercial assays** |  |  |
| Mitochondrial Stress Test Kit | Agilent | 103015-100 |
| XF DMEM Medium | Agilent | 103575-100 |
| **Experimental models: Organisms/strains** |  |  |
| C57BL/6J *Mus Musculus* – 6 weeks of age | Jackson Laboratories | RRID:<br>IMSR_JAX:000664 |

### Resource availability

#### Lead contact

Further information and requests for resources and reagents should be directed to and will be fulfilled by the lead contact, Kimberly Dunham-Snary.

#### Data and code availability

- Data reported in this article is available upon request from the lead contact.
- This paper does not report original code.
- Any additional information required to reanalyze the data reported in this work paper is available from the lead contact upon request.

### Experimental model and study participant details

#### Animals

All animal studies carried out were approved by the University Animal Care Committee at Queen’s University (Protocol 2023-2390). Briefly, 4-5-week-old male C57BL/6J mice (n = 30) were purchased from Jackson Laboratories (JAX) and fed standard rodent diet (5020 – Mouse Diet 9F; LabDiet, St. Louis, MO, USA) and water *ad libitum* for 1-2 weeks until 6-weeks of age; six weeks was chosen to ensure minimal age-associated changes in metabolism, redox balance, and inflammation^29^. Animals were anesthetized with 3-5% isoflurane, subsequently euthanized via cardiac puncture, and cervical dislocation as a secondary means of euthanasia.

### Method details

#### Tissue Collection

Animals were anesthetized via induction with isoflurane and blood was collected via cardiac puncture into a syringe pre-filled with 0.1 mL 3.2% sodium citrate for isolation of platelets. Gastrocnemius muscle was excised, and tissue was immediately placed in ice-cold BIOPS buffer (2.77 mM CaK_2_EGTA, 0.5 mM Dithiothreitol, 20 mM Imidazole, 7.23 mM K_2_EGTA, 50mM MES Hydrate, 6.56 mM MgCl_2_×6H2O, 5.77 mM Na_2_ATP, 15 mM Na_2_Phosphocreatine, 20 mM Taurine; pH 7.2) for respirometry.

#### Platelet Isolation

Whole blood was centrifuged at 250 x g for 2 min at room temperature (RT). Platelet rich plasma (PRP) was collected. Erythrocyte lysis buffer supplemented with PGI_2_ (0.02 mg/mL; “platelet buffer”) was added to whole blood, which was then centrifuged at 250 x g for 2 min RT. PRP was collected and combined with the previous fraction to maximize platelet cell counts. PRP was centrifuged at 400 x g for 5 min RT and the supernatant (platelet poor plasma) was discarded. The platelet pellet was resuspended in platelet buffer and incubated at RT for 6 minutes to remove remaining red blood cell contamination before centrifugation for 5 min at 400 x g RT. The supernatant was discarded, and the platelet pellet was resuspended in 50 µL platelet buffer.

Platelet count was obtained via Beckman Coulter Counter Z Series, using a 50 μm aperture with the upper and lower limits set to 5.1 fl and 4.3 fl, respectively, to account for the size difference between mouse and human platelets^30^. Each sample was measured in triplicate and averaged to obtain a final cell concentration.

#### Flow Cytometry

Platelet samples were isolated from 3 mice, with half of each sample receiving 1 U/ml thrombin at 37 °C for 15 min. Platelets were washed with Flow Cytometry Staining (FACS) buffer (800 g, 5 min, and 21°C), and each sample was incubated with CD41-APC (platelet marker), CD45-AlexaFluor700 (total leukocyte marker), and P-Selectin-PE (platelet activation marker) antibodies in FACS buffer for 30 min at 4°C. Samples were washed twice with FACS buffer, resuspended in FACS buffer, and analyzed using a SH800 Cell Sorter (Sony Biotechnology). Flow cytometry confirmed that pure (**Fig. S1**) and quiescent (**Fig. S2**) platelet isolations were used for respirometry. Fluorescence minus one (FMO) controls were used to determine gating and staining quality.

#### Murine Platelet Mitochondrial Stress Test (XFe24)

Platelet mitochondrial oxygen consumption rate (OCR) was measured via high-resolution respirometry (HRR) using a Seahorse Extracellular Flux Analyzer (XFe24; Agilent). Cell seeding density, oligomycin titration, and FCCP titration were performed for XFe24 assay optimization prior to study commencement. Platelets were seeded at a density of 25 x 10^6^, but preliminary data analysis showed oxygen (O_2_) levels were approaching 0 mmHg during measurement cycles, so to avoid inducing hypoxic conditions, seeding density was reduced to 20 x 10^6^ platelets per well. All platelet OCR measurements were normalized to cell seeding density. Each platelet sample was run in duplicate if enough platelets were available. The plate was centrifuged at 250 x g for 1 min in a swing bucket centrifuge (acceleration = 1, zero brake for deceleration), and then rotated and centrifuged again under the same conditions to ensure even coating of platelets in each well. Platelet buffer was aspirated and Seahorse XF DMEM Medium (5mM HEPES, 15 mM glucose, 2 mM glutamine, 1 mM pyruvate; pH 7.45) was added to each well.

After baseline respiration measurements (*basal*), respiration was measured during sequential injections of oligomycin (2.5 µM), FCCP (0.4 µM), and antimycin A/rotenone (5µM each). Injection of antimycin A/rotenone inhibits ETC complexes III and I, respectively, revealing *non-mitochondrial* OCR, which is subtracted from all other respiration parameters. Injection of oligomycin inhibits ATP synthase, revealing respiration attributed to *proton leak*, and when the minimum OCR after oligomycin injection is subtracted from *basal respiration*, respiration linked to ATP production is revealed (*ATP-linked*). Injection of FCCP uncouples the ETC causing respiration to reach its *maximal* OCR. Subtracting *basal* from *maximal* respiration reveals *reserve capacity* (ResCap), or the capacity of the sample to respond to conditions of high energy demand. *Coupling Efficiency (CE (%))* was calculated as ATP-linked/basal x 100. One sample had ATP-linked, proton leak, and CE (%) data excluded due to technical artifact upon oligomycin injection (e.g., injection if an air bubble).

#### Permeabilized Muscle Fibers

White gastrocnemius muscle tissue was placed in ice-cold BIOPS (2.77 mM CaK_2_EGTA, 0.5 mM Dithiothreitol, 20 mM Imidazole, 7.23 mM K_2_EGTA, 50mM MES Hydrate, 6.56 mM MgCl_2_×6H_2_O, 5.77 mM Na_2_ATP, 15 mM Na_2_Phosphocreatine, 20 mM Taurine; pH 7.2). All visible remnant connective tissue and fat was removed from the muscle sample prior to, and during, mechanical separation. Muscle bundles were formed by gently separating the fibers along the longitudinal axis of the muscle using needle-tipped forceps under a microscope (AmScope, USA). Fiber bundles were weighed in ~1.5 mL of tared ice-cold BIOPS while obtaining fiber bundle wet weights to ensure that fibers remained relaxed and hydrated. Bundle wet weights were obtained in duplicate and the average of the two values was used to normalize rates of mitochondrial respiration to fiber bundle mass. Fiber bundles remained in ice cold BIOPS until chemical permeabilization with the cholesterol-specific detergent saponin. Muscle bundles were then chemically permeabilized for 30 minutes in BIOPS with 40 μg/mL saponin at 4°C. Following permeabilization, bundles were washed in Buffer Z (5 mg/mL fatty acid-free BSA, 1 mM EGTA, 30 mM KCl, 10 mM KH_2_PO_4_, 105 mM K-MES, 5 mM MgCl_2_×6H_2_O; pH 7.2) for 15 min at 4°C.

#### Murine Skeletal Muscle Respirometry (O2k)

Mitochondrial OCR was measured via HRR using the Oxygraph 2k (O2k; Oroboros Instruments, Innsbruck Austria). All mitochondrial respiration measurements were obtained while the chamber oxygen concentration was between 400-180 μM. All respiration measurements were performed in 2 mL of Buffer Z, with stirring at 750 r.p.m and 37°C. Respiration measurements were performed in duplicate in the presence of the myosin ATPase inhibitor blebbistatin (5μM), which was added to each chamber prior to bundle insertion and chamber oxygenation^31^. Data was acquired every 2 seconds with the uncorrected rate of mitochondrial oxygen consumption (pmol/s/mL) calculated from 40 data points. Raw oxygen consumption values were reported in pmol/s/mL on DatLab and data corrected for bundle wet weight (wt.) are shown in pmol/s/mg wet wt. Substrate addition occurred as follows: pyruvate (5mM), malate (2 mM), glutamate (10 mM), ADP titrations (500 µM, 5 mM, 15 mM), cytochrome c (10 µM), succinate (20 mM), oligomycin (2.5 µM), FCCP titration (0.5 µM, 1.0 µM, 1.5 µM, 2.0 µM), antimycin A (2.5 µM)/rotenone (0.5 µM).

All parameters of respiration derived from these measurements were corrected for background fiber bundle respiration in the absence of exogenous substrates and had non-mitochondrial oxygen consumption (*Rox*), induced by antimycin A and rotenone addition, subtracted out to obtain true mitochondrial respiration values. *OXPHOS capacity* (P) was measured after addition of succinate; *ET capacity* (E) was determined from the maximal respiration rate after FCCP titration – these values were later excluded due to the documented phenomenon that E ≤ P in murine skeletal muscle^32,33^; *LEAK* (L) was defined as respiration after oligomycin addition; *ATP-linked* respiration was calculated by subtracting L from P; *CE (%)* was calculated as ATP-linked/P x 100; *respiratory control ratio* (RCR) was calculated as P/L^34,35^.

Permeabilized muscle fibers from the same gastrocnemius sample were run in duplicate; n=12 samples had data from only one muscle bundle included in analysis due to negative fiber bundle background respiration values following correction for instrumental background and/or low responsiveness to ADP titration *i.e.,* bundles that did not respire in response to ADP. 6 muscle samples had LEAK, ATP-linked, and CE (%) excluded from analysis due to erroneous addition of FCCP before oligomycin prior to a protocol change; 5 samples were excluded from analysis due to increased respiration in response to cytochrome c addition, indicating compromised mitochondrial membrane integrity^36^. ET-capacity and ResCap measurements were excluded from correlation analysis due to the observation that FCCP did not induce maximal respiration rates, in line with previous literature on murine skeletal muscle respirometry^32,33^.

### Quantification and statistical analysis

#### Statistics

All statistical analyses were performed in GraphPad Prism. Respirometry parameters from the XFe24 and O2k were input into a table, and each parameter was assessed for normality using a Shapiro-Wilks test. XF maximal is the only respiratory parameter that failed the normality test. Outliers were removed if they were outside 3 standard deviations from the mean of each parameter (1 point from OXPHOS, 1 point from XF CE (%), 1 point from XF ResCap (%)). Correlations were performed to compute Spearman ρ (non-normal data) or Pearson r values (normally distributed data), using a two-tailed p-value with confidence intervals set to 95%. Correlations were considered significant if p < 0.05.

## Results

### Murine Skeletal Muscle High Resolution Respirometry (HRR)

Murine skeletal muscle respirometry in white gastrocnemius muscle was performed using the Oxygraph 2k (O2k; Oroboros), performing a substrate-uncoupler-inhibitor-titration (SUIT) protocol that mimics the Agilent Mitochondrial Stress Test assay (**Fig. 1A**), but FCCP measurements were excluded (see methods section) in accordance with previous literature on murine skeletal muscle respirometry^32,33^. Mean and SD values for muscle respiration are displayed in **Figure 1B**.

**Figure 1.**
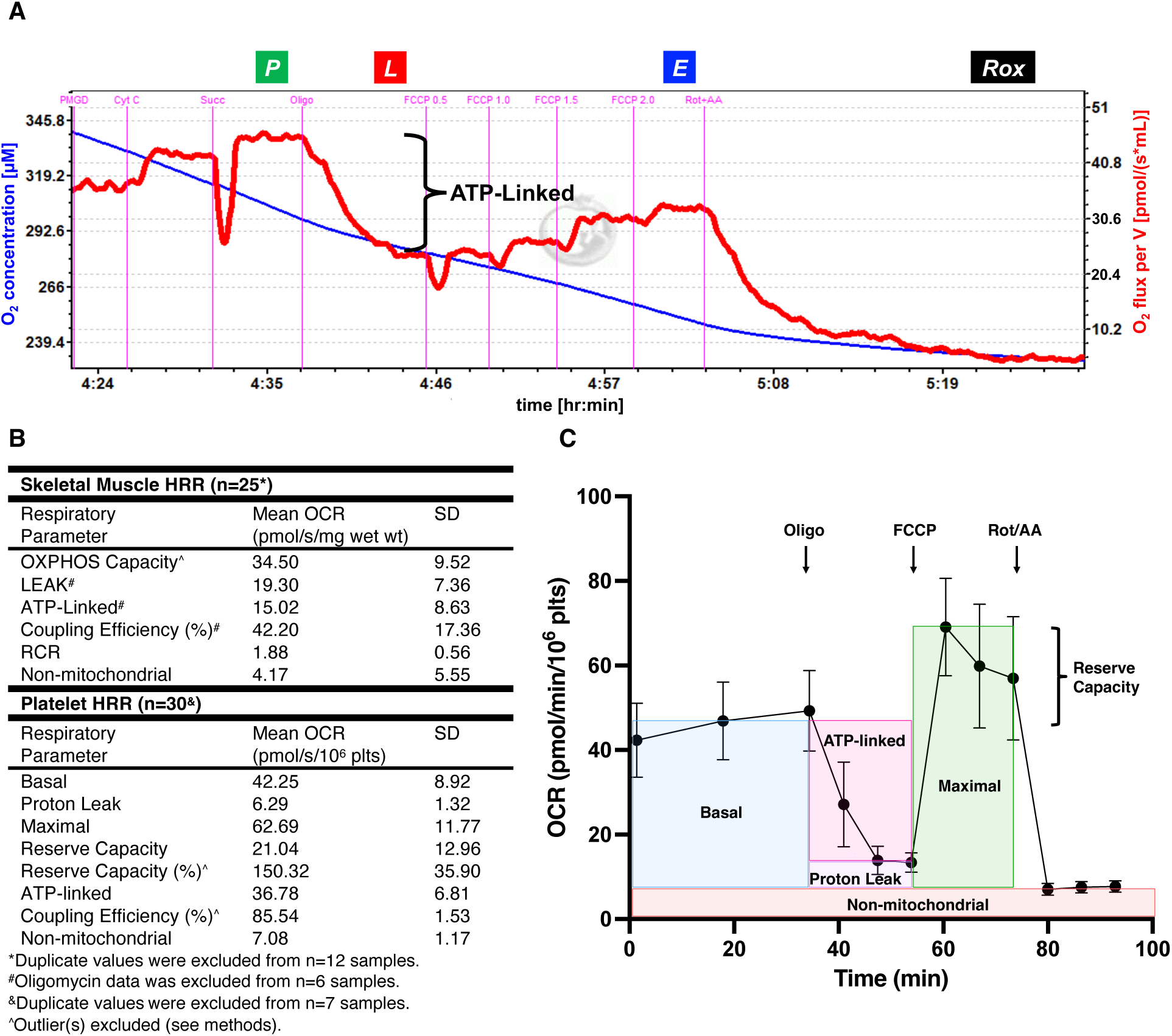
Summary of HRR data from skeletal muscle and platelets. **A. Representative O2k trace of permeabilized murine gastrocnemius muscle respirometry (n=1).** Red and blue lines indicate O_2_ flux and O_2_ concentration in chamber, respectively. Bracket indicates portion of respiration liked to ATP-production (P–L). Pink lines indicate substrate/effector addition. **B. Table summarizing mean OCR values for muscle and platelet HRR parameters. C. Platelet bioenergetics from healthy male 6-week-old C57BL/6J mice (n=30).** Error bars represent SD. Effector injections indicated by black arrows. Respiration parameters are indicated on the trace. *P – pyruvate; M – malate; G – Glutamate; D – ADP; Cyt C – cytochrome C; Succ – succinate; P – OXPHOS capacity; L – LEAK; E – ET capacity; Rox – residual oxygen consumption; Oligo – Oligomycin; FCCP - Carbonyl cyanide-p-trifluoromethoxyphenylhydrazone; Rot/AA – Rotenone/Antimycin A; RCR – Respiratory control ratio (OXPHOS/LEAK)*.

### Murine Platelet HRR

We have optimized a protocol for executing the Agilent Mitochondrial Stress Test in murine platelets using the Seahorse Extracellular Flux Analyzer (XFe24; see methods section). Mean and SD values for platelet respiration are displayed in **Figure 1B**. Grouped kinetic data from the XFe24 is shown in **Figure 1C**.

### Correlation Analysis

Mitochondrial respiration parameters obtained from platelets via the XFe24 include basal, proton leak, ATP-linked, maximal, reserve capacity (ResCap), reserve capacity as % above basal (ResCap (%)), coupling efficiency (CE (%)), and non-mitochondrial respiration. CE (%) contextualizes ATP-linked respiration as a proportion of basal respiration. Mitochondrial respiration parameters obtained from skeletal muscle via the O2k include OXPHOS, LEAK, ATP-linked, CE (%), respiratory control ratio (RCR), and non-mitochondrial respiration. Respiration values from both tissues are presented in a correlation matrix (**Fig. 2**).

**Figure 2.**
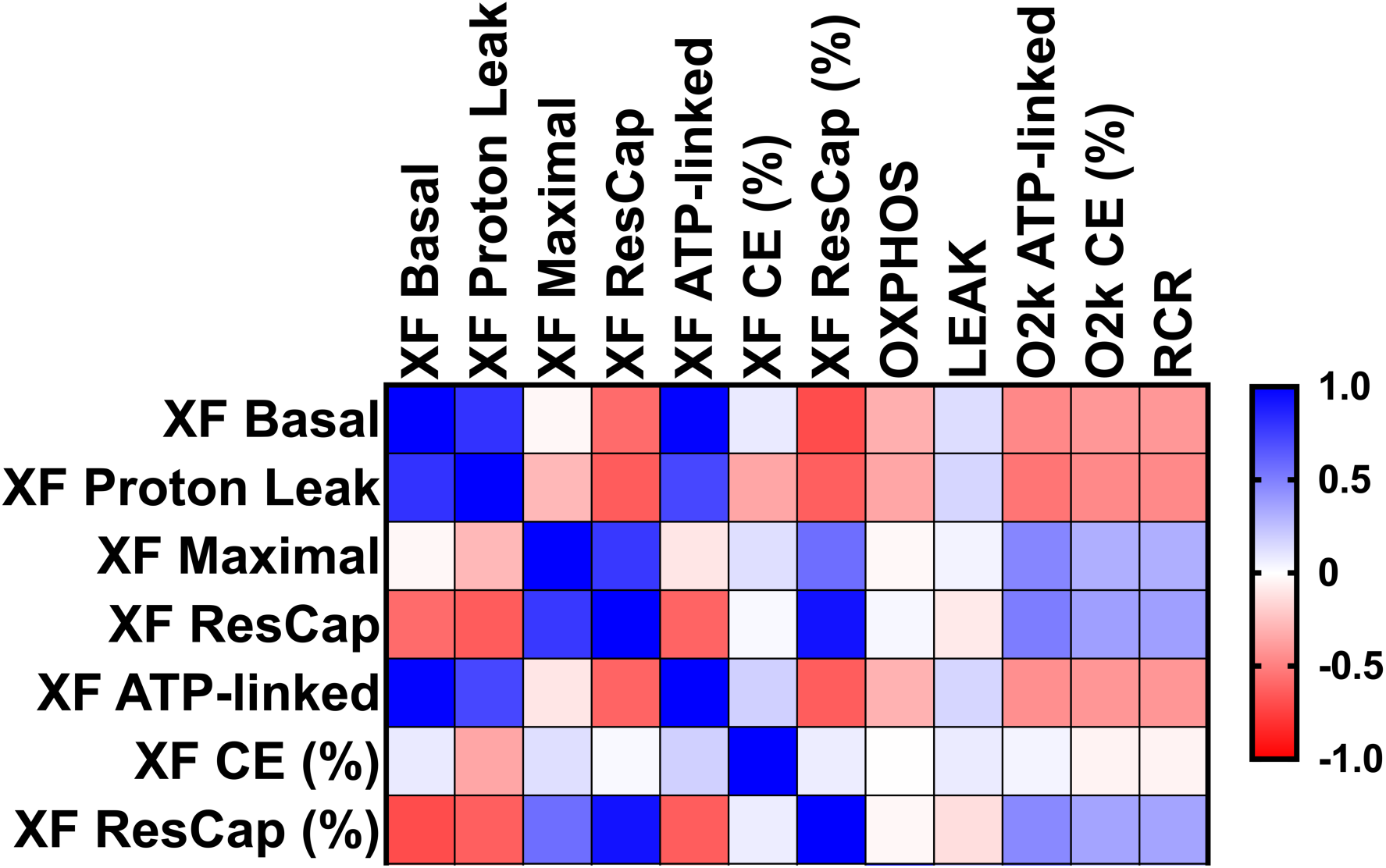
Spearman correlation matrix performed on XFe24 (platelet) and O2k (muscle) measurements. Blue and red represent positive and negative Spearman r values, respectively. Saturation indicates the strength of correlation. *ResCap – Reserve Capacity; CE (%) – % Coupling Efficiency; RCR – Respiratory Control Ratio*.

### Correlation of Platelet and Muscle Respirometry

We have discovered multiple physiologically relevant correlations between platelet and skeletal muscle bioenergetics in healthy male mice, providing significant insights into mitochondrial coupling. In a context-dependent manner, mitochondrial (un)coupling can support biologically protective processes including buffering during ROS-favouring conditions or can also indicate global mitochondrial dysfunction^37^.

Platelet basal respiration negatively correlates with skeletal muscle ATP-linked respiration (**Fig. 3A**) and RCR (**Fig. 3B**), demonstrating concomitant, opposing changes between platelet baseline and skeletal muscle ATP-linked respiration. RCR, an index of how coupled respiration is to ADP phosphorylation^34,35^, negatively correlates with platelet proton leak (**Fig. 3D**). Since higher and lower RCR values indicate coupled organelles and higher LEAK respiration, respectively, *the negative correlation between muscle RCR and platelet proton leak link the coupling states of these tissues*.

**Figure 3.**
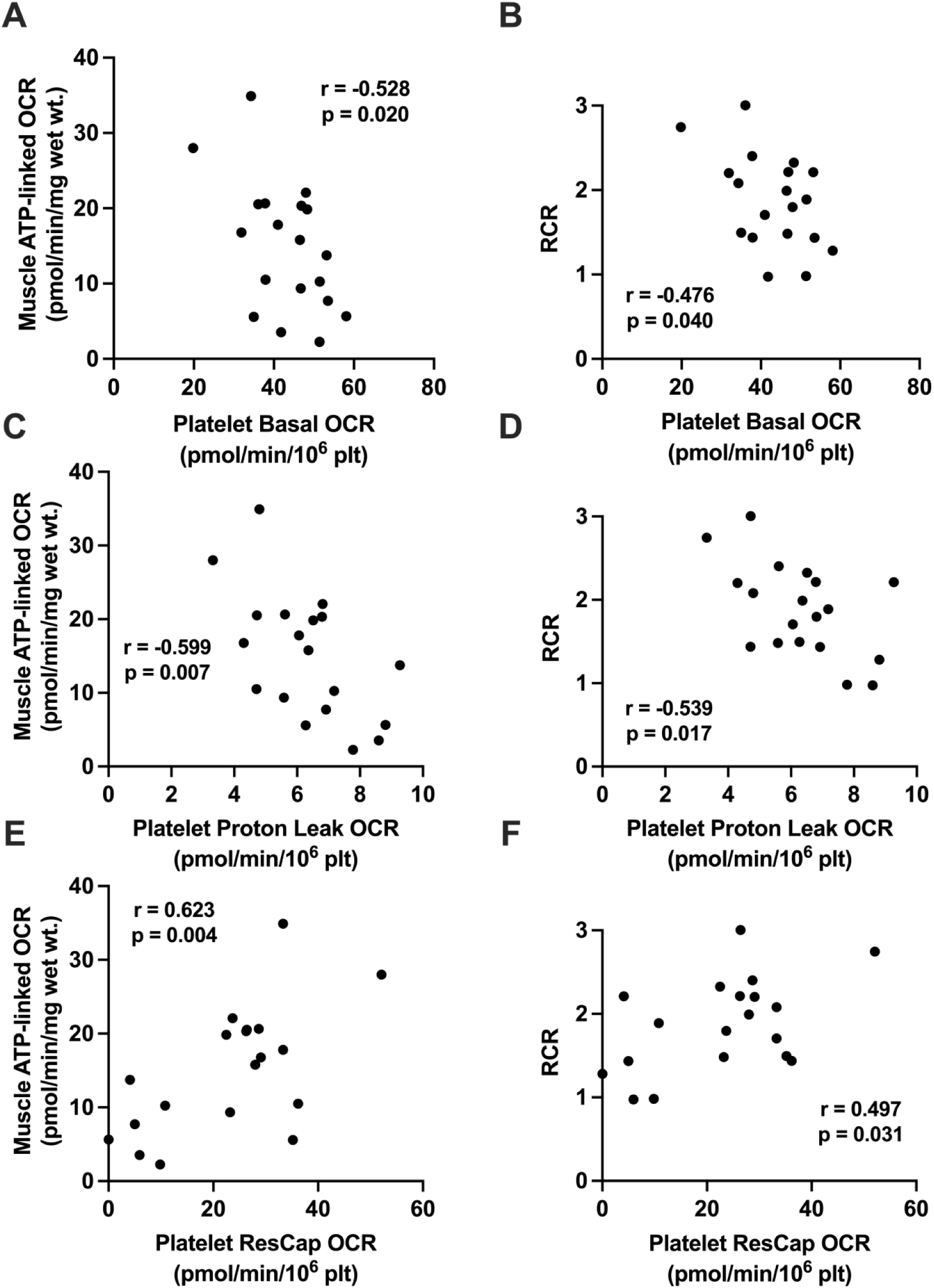
Pearson correlations between platelet and skeletal muscle HRR. Skeletal muscle (y-axes) and platelet (x-axes) respiration data were normalized to tissue wet weight (wt.) and cell seeding density, respectively. Pearson r and p values are reported on each graph. **A.** Platelet basal and muscle ATP-linked respiration. **B.** Platelet basal and muscle RCR. **C.** Platelet proton leak and muscle ATP-linked respiration. **D.** Platelet proton leak and muscle RCR. **E.** Platelet ResCap and muscle ATP-linked respiration. **F.** Platelet ResCap and muscle RCR.

Platelet proton leak also negatively correlates with muscle ATP-linked oxygen consumption (**Fig. 3C**) and CE (%, r = −0.533, p = 0.019), demonstrating the ability of platelet respiration as a circulating record of muscle mitochondria coupling state. Furthermore, platelet maximal respiration correlates positively with muscle ATP-linked respiration (ρ= 0.472, p = 0.041), and correlation was strengthened when corrected for variation in basal respiration by assessing reserve capacity (ResCap). ResCap correlates positively with muscle ATP-linked respiration (**Fig. 3E**) and RCR (**Fig. 3F**). When expressed as a percentage of baseline (maximal/basal x 100), platelet ResCap (%) and muscle ATP-linked still positively correlate (r = 0.490, p = 0.039). *These relationships further link the respiratory capacity of platelets with the coupling state of the skeletal muscle*.

## Discussion

Correlations with skeletal muscle ATP-linked and RCR (**Fig. 3**) indicate that platelet bioenergetics are a reliable circulating biomarker of skeletal muscle coupling state in healthy mice. As platelet maximal respiration and ResCap increase, we have demonstrated that ATP-linked respiration in skeletal muscle also increases; skeletal muscle mitochondria are operating in a more coupled, energy-efficient state. Therefore, when platelets exhibit greater maximal respiration and an augmented capacity to increase respiration in response to energy demand, the skeletal muscle is more ‘fuel-efficient’.

This is the first report assessing the correlation between platelet and skeletal muscle mitochondrial bioenergetics in mice, and the first robust demonstration of the utility of platelet bioenergetics as a surrogate for skeletal muscle metabolism in mice. The correlations demonstrated here in healthy mice set the stage for investigating how platelet metabolism responds to nutrient overload and genetic factors linked to CMD susceptibility and pathogenesis. This study is limited to the use of a homogenous, healthy cohort of mice; future work should explore these correlations in an established disease model of CMD. While numerous analyses of muscle bioenergetics in animal models of obesity and diabetes are documented, results are conflicting. Adult Zucker rats exhibit impaired respiration in mitochondria isolated from soleus muscle that is not present in young rats^38^. Conversely, obese Wistar rats exhibit increased OXPHOS and ET capacities in diaphragm muscle compared to lean controls^39^. In mitochondria isolated from cardiac muscle of obese C57Bl/6J mice, Boardman and colleagues demonstrated decreased pyruvate- and palmitate-stimulated state 3 and state 4 respiration compared to age-matched controls, with no change in RCR^40^. As RCR is calculated by state 3/state 4 (OXPHOS/LEAK), the decrease in OXPHOS without a change in RCR is due to the concomitant decrease in state 4 respiration. Similar results were demonstrated in isolated cardiac mitochondria of obese Zucker rats, exhibiting decreased state 3 and state 3u respiration compared to lean controls^41^.

When assessing skeletal muscle in animal model of diabetes, respiration trends from extensor digitorum versus soleus muscle show conflicting patterns; LEAK, CI-linked OXPHOS, CI+CII-linked OXPHOS, and ET capacity are increased in extensor digitorum muscle of *db/db* mice, but CI-linked OXPHOS is decreased in soleus muscle^42^, supporting the finding that diabetes differentially impacts mitochondrial metabolism in skeletal muscle subtypes^43^. An interesting avenue of further investigation is assessing the applicability of platelet bioenergetics as compared to a mixed fiber type (e.g., vastus lateralis) or predominantly type I, “slow twitch” fiber muscle (e.g., soleus) metabolism, as the work herein was performed in white gastrocnemius muscle (predominantly type II, “fast twitch” fiber). When comparing tibialis anterior muscle of lean non-diabetic *fa*/+ and obese diabetic *fa*/*fa* adult male Zucker diabetic fatty rats, Wessels et al. found no differences in CI- or CII-linked OXHPOS, but increased CI-linked RCR in diabetic animals suggesting uncoupling^44^.

We have demonstrated that platelet proton leak negatively correlates with muscle ATP-linked (**Fig. 3A**) and RCR (**Fig. 3B**) and conclude that the coupling state of platelets and skeletal muscle are related in healthy mice; when platelets have a higher proton leak (less coupled), we show the skeletal muscle has a lower ATP-linked respiration and RCR (i.e., are also less coupled). Therefore, *we propose that the coupling state of platelet mitochondria reflects the coupling state of muscle mitochondria in the same animal (****Figure 4***).

**Figure 4.**
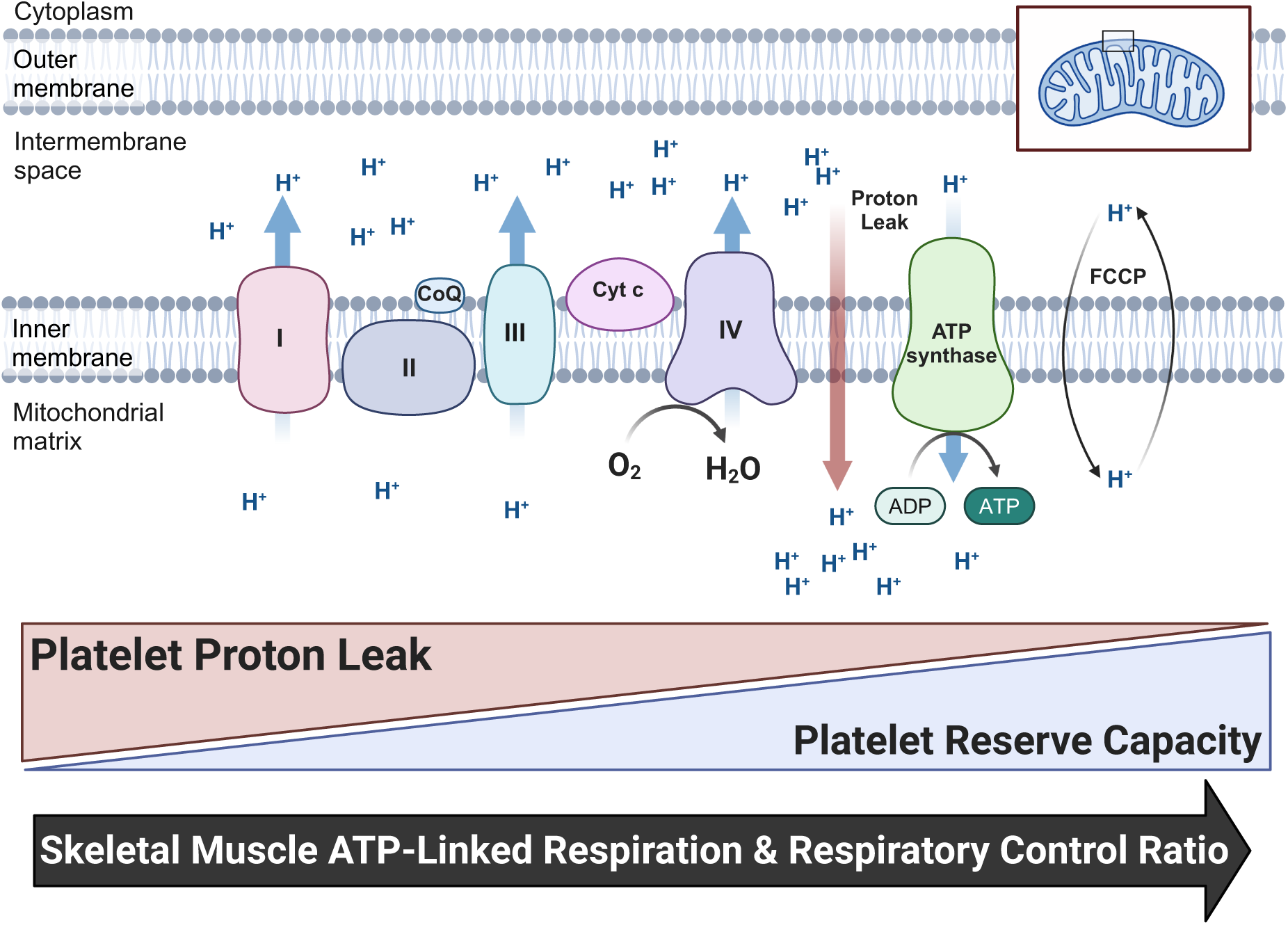
Proposed model of platelet mitochondrial respiration as a liquid biopsy of skeletal muscle mitochondrial health. ETC complexes pump protons in the intermembrane space creating an electrochemical gradient driving conversion of ADP to ATP via ATP synthase (green). Some protons ‘leak’ back into the matrix (red arrow) and are uncoupled from ATP-production. FCCP allows protons to move freely across the membrane (black arrows), uncoupling respiration and causing OCR at complex IV (purple) to reach its maximum. *CoQ – Coenzyme Q; Cyt c – cytochrome c*.

In human obesity, Vijgen and colleagues have shown that skeletal muscle of morbidly obese patients has decreased CI+CII-linked OXPHOS capacity, with unchanged LEAK respiration compared to lean controls^45^. Further, they found that weight loss induced one year after bariatric surgery diminished the contribution of LEAK to OXPHOS. As well, Phelix et al. demonstrated that obese individuals have decreases in CI- and CI+CII-linked OXPHOS, as well as ET capacity compared to lean controls^46^. These findings contrast work from Ara et al., who reported that obesity did not induce changes in skeletal muscle mitochondrial respiration compared to lean or post-obese controls^47^; they report a trending increase in CI-linked OXPHOS in muscle from obese participants after respiration data was normalized to citrate synthase activity. Similarly, when investigating the impact of “pre-diabetes” on skeletal muscle respiration, Szczerbinski et al. found no change in any mitochondrial respiration parameters before or after exercise intervention^48^. These findings are somewhat incongruent with the current understanding of skeletal muscle metabolic adaptations to exercise^49^, therefore, further investigation into how the “pre-diabetes state” influences skeletal muscle mitochondrial respiration is warranted. It is well documented that diabetes confers a deficit in skeletal muscle respiration^44,46,50–54^; an inverse relationship between HbA1c and mitochondrial respiration parameters in diabetic and non-diabetic individuals has been demonstrated^50,53^.

Human studies investigating platelet bioenergetics have yielded promising early results, but *correlation between skeletal muscle and platelet mitochondrial respiration has yet to be assessed in CMD patients*. Platelets from type II diabetic patients exhibit decreased basal and ATP-linked OCR compared to age-matched controls, and this decrease was associated with markers of oxidative stress^55^. When comparing platelet and muscle bioenergetics of healthy women of various BMI status, Rose and colleagues report that CI-linked LEAK respiration and OXPHOS coupling efficiency both positively correlate between the tissues^24^. As well, Braganza et al. showed correlation between muscle and platelet bioenergetics in young versus aged individuals^23^. However, Westerlund et al. found few significant correlations between platelet and skeletal muscle metabolism in athlete and non-athlete controls; they revealed negative correlations in background respiration and NS-OXPHOS capacity^56^. They also demonstrated that augmented mass-specific muscle respiration shown in athletes was not reflected in platelet metabolism, but in a subset of patients with primary mitochondrial disease there were significantly altered platelet bioenergetics relative to healthy controls. In animal model, Tyrell demonstrated correlation between platelet and skeletal muscle in African green monkeys^25^. More work is necessary to explore the potential of platelet bioenergetics as a muscle-specific mitochondrial biomarker.

The work described herein establishes novel correlations between platelet and skeletal muscle bioenergetics in healthy mice, demonstrating that platelet bioenergetics can serve as a surrogate for skeletal muscle mitochondrial health. These findings provide a foundation for future work investigating how platelet bioenergetics respond to metabolic challenges, such as nutrient overload or genetic factors linked to CMD susceptibility.

## Conclusion

Platelets have the potential to serve as a muscle-specific bioenergetic marker and this proof-of-concept study lays the groundwork for studying the impact of other genetic, epigenetic, and environmental factors on both platelet and muscle bioenergetics in animal models, as well as for future human studies in healthy individuals and CMD patients. While this study was limited by lack of an experimental disease model, we demonstrated strong correlation between the measures of interest in healthy mice, warranting further investigation into this marker in the context of CMD. If platelet mitochondria can reflect early metabolic changes associated with CMD, especially *before* perturbations in traditional markers such as fasting blood glucose or HbA1c, this method may serve as a potent clinical tool for assessing CMD risk and implementing prevention and intervention strategies before the onset of overt disease.

## Acknowledgments

The authors would like to acknowledge Benjamin Pieter Ott for guidance with statistical analysis. We would also like to acknowledge Dr. M. Ryan Smith (Emory University) and Mr. Jeffrey Mewburn (Queen’s University) for providing guidance with platelet isolation, counting, and plating.

## Funding

Canadian Institutes of Health Research (PJT190103) KDS

Tier II Canada Research Chair (CRC-2020-00192) KDS

Canada Foundation for Innovation – John Evans Leaders Fund (41511) KDS

Queen’s University Faculty of Health Sciences (6032495) KDS

Banting Research Foundation and Mitacs Discovery Award (6035577) KDS

Department of Medicine, School of Medicine, Queen’s University (6034430) KDS

Natural Sciences and Engineering Research Council of Canada (NSERC-00104-2020) CM

## Author contributions

Conceptualization: MSW, CM, KJDS

Methodology: MSW, EJF, PDAL

Investigation: MSW, EJF, JB, JLMHV, PDAL

Formal Analysis: MSW

Resources: CM, KJDS

Visualization: MSW, PDAL

Funding acquisition: KJDS

Supervision: CM, KJDS

Writing – original draft: MSW, EJF

Writing – review & editing: KJDS

## Competing interests

Authors declare that they have no competing interests.

## Data and materials availability

All data are available in the main text or the supplementary materials. Raw data files are available from the corresponding author upon request.

## Supplemental Information

Document S1. Figures S1-S2.

## Notes

### Competing Interest Statement

The authors have declared no competing interest.

