## Supplementary Figures for "Platelet bioenergetics correlate with skeletal muscle metabolism in C57BL/6J mice"

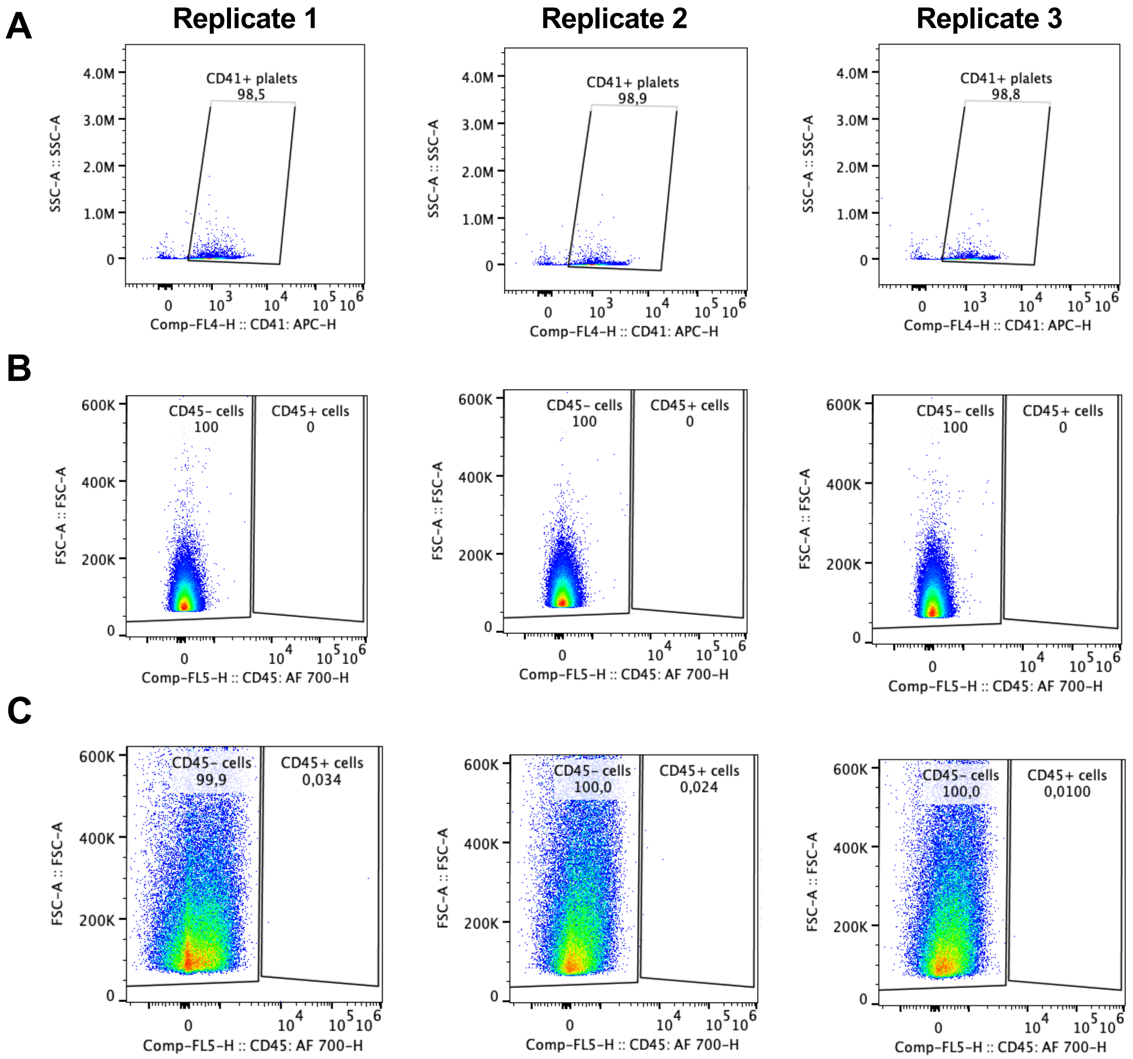


**Figure S1. Assessment of platelet isolation purity. A.** CD41 positive cells (platelets) from three murine samples demonstrating the purity of the platelet isolates. Percentage of total events is indicated on each graph with decimals denoted with commas. **B-C.** CD45 (leukocyte marker) negative (left) and positive (right) events in quiescent (**B**) and thrombin-activated (**C**) platelet isolates demonstrating purity of the platelet samples. Percentage of total events is indicated on each graph with decimals denoted with commas.

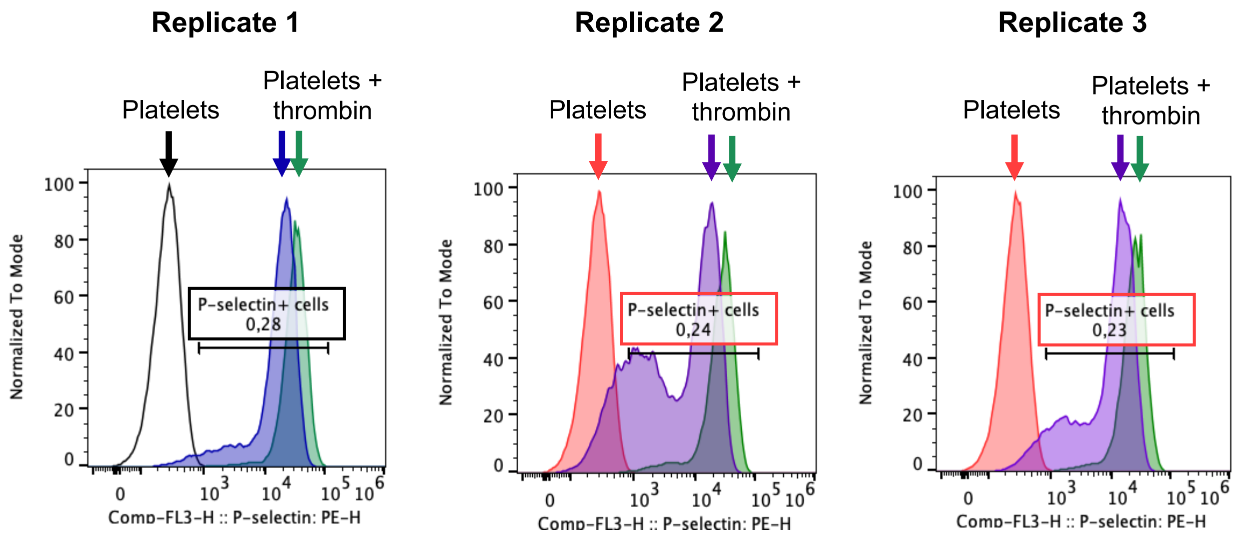


**Figure S2. Assessment of platelet isolation activation status.** P-selectin positive cells (activated platelets) in unstimulated platelet isolates and platelets incubated with thrombin (a platelet activator) demonstrating that the platelet isolation protocol itself does not activate platelets. Percentage of total events for the quiescent samples is indicated on each graph with decimals denoted with commas.
